# A tubulin-binding protein that preferentially binds to GDP-tubulin and promotes GTP exchange

**DOI:** 10.1101/2023.05.09.539990

**Authors:** Wesley J. Yon, Taekjip Ha, Yixian Zheng, Ross T.A. Pedersen

## Abstract

α- and β-tubulin form heterodimers, with GTPase activity, that assemble into microtubules. Like other GTPases, the nucleotide-bound state of tubulin heterodimers controls whether the molecules are in a biologically active or inactive state. While α-tubulin in the heterodimer is constitutively bound to GTP, β-tubulin can be bound to either GDP (GDP-tubulin) or GTP (GTP-tubulin). GTP-tubulin hydrolyzes its GTP to GDP following assembly into a microtubule and, upon disassembly, must exchange its bound GDP for GTP to participate in subsequent microtubule polymerization. Tubulin dimers have been shown to exhibit rapid intrinsic nucleotide exchange in vitro, leading to a commonly accepted belief that a tubulin guanine nucleotide exchange factor (GEF) may be unnecessary in cells. Here, we use quantitative binding assays to show that BuGZ, a spindle assembly factor, binds tightly to GDP-tubulin, less tightly to GTP-tubulin, and weakly to microtubules. We further show that BuGZ promotes the incorporation of GTP into tubulin using a nucleotide exchange assay. The discovery of a tubulin GEF suggests a mechanism that may aid rapid microtubule assembly dynamics in cells.

## Introduction

Microtubules represent one type of the eukaryotic cytoskeleton required for key cellular functions including transportation of cargos via motor proteins, relaying of mechanical signals to the interphase nucleus, and proper segregation of chromosomes during cell division (Gudimchuk and McIntosh, 2021; Reck-Peterson et al., 2018; Zheng, 2010; Kirby and Lammerding, 2018). Microtubules are assembled from tubulin, a heterodimer of α- and β-tubulin. α-tubulin is constitutively bound to guanosine triphosphate (GTP) at its non-hydrolyzing and non-exchangeable “N-site”, whereas β-tubulin binds to and rapidly exchanges both GTP and GDP at its exchangeable “E-site” where GTP can be hydrolyzed into GDP (Jacobs et al., 1974; Brylawski and Caplow, 1983). Tubulin heterodimers with GTP-bound β-tubulin (hereinafter referred to as GTP-tubulin) polymerize into microtubules (Desai and Mitchison, 1997). As the microtubule elongates, longitudinal and lateral interactions of incorporated tubulin dimers activate β-tubulin GTPase activity, stochastically hydrolyzing the bound GTP and forming GDP-tubulin in the microtubule lattice (Shemesh et al., 2023; Nogales et al., 1998a). As a result, the growing ends of microtubules contain GTP-tubulin, called the GTP cap. GTP hydrolysis induces tubulin dimer conformational changes which cause destabilization of the microtubule lattice (Alushin et al., 2014). In the absence of the GTP cap, microtubules transition from growth to shrinkage, termed catastrophe (Horio and Murata, 2014). Microtubule disassembly releases free GDP-tubulin, which must exchange its bound GDP for GTP before it can take part in microtubule polymerization again.

GTPases generally exhibit slow intrinsic rates of nucleotide exchange (t_1/2_ > 30 min) and require <u>g</u>uanine nucleotide <u>e</u>xchange factors (GEF) to accelerate nucleotide dissociation by orders of magnitude for proper biological function (Bischoff and Ponstingl, 1991; Blaise et al., 2021; Marshall et al., 2012; Self and Hall, 1995; Chardin et al., 1993; Killoran and Smith, 2019). Many GEFs have been characterized for different GTPase families, and GEFs within a family share common catalytic domains such as the CDC25 domain for RasGEFs and the Sec7 domain for ArfGEFs (Bos et al., 2007). GEFs between different GTPase families do not share homologous protein sequences making it difficult to identify novel GEFs from primary structure alone (Koch et al., 2016). Despite the lack of sequence similarity, studies have shown a common mechanism by which GEFs mediate the release of GTPase-bound nucleotides by destabilizing the magnesium ion in the nucleotide binding pocket required for stable nucleotide binding (Vetter and Wittinghofer, 2001; Béraud-Dufour et al., 1998). GEFs bind to GTPases without specificity to their nucleotide states and mediate the exchange of either bound GTP or GDP with nucleotides in solution (Boor et al., 2015; Bos et al., 2007; Bischoff and Ponstingl, 1991). In cells, the 10:1 ratio of intracellular GTP:GDP ensures that the GTP-bound form is the dominant GTPase species when acted upon by a GEF (Traut, 1994). In contrast to slow measured rates of nucleotide exchange of many GTPases, the *in vitro* dissociation of GDP from β-tubulin is rapid (t_1/2_ = 5 s) (Brylawski and Caplow, 1983). Therefore, it is believed that free tubulin in the cell is able to quickly exchange into the GTP-bound form leading to the view that a tubulin GEF is not needed. However, there is an 18-fold increase in microtubule turnover in mitosis compared to in interphase, representing a considerable increase in demand for GTP-tubulin (Saxton et al., 1984). Additionally, microtubule dynamics have been shown to be diffusion-limited in metaphase *Xenopus* egg extracts, and energy consumption outpaces energy production during cell division (Galichon et al., 2024; Geisterfer et al., 2020). In this context, the more rapid polymerization and depolymerization of microtubules may necessitate tubulin GEF activity to sustain elevated microtubule turnover during spindle assembly.

BuGZ (<u>Bu</u>b3-interacting and <u>G</u>LEBS motif-containing protein <u>Z</u>NF207) is a recently discovered protein that binds to tubulin and microtubules (Jiang et al., 2015). In mitosis, BuGZ regulates spindle assembly by promoting microtubule polymerization and proper microtubule-kinetochore attachment by binding and stabilizing Bub3, a spindle assembly checkpoint protein. As a result, reduction of BuGZ results in prometaphase arrest with defects in spindle morphology and chromosome alignment (Jiang et al., 2014; Toledo et al., 2014). BuGZ is evolutionarily conserved in animals and plants, and the plant BuGZ may also regulate microtubule-kinetochore interactions as it contains a predicted Bub3 binding domain (Chin et al., 2022). BuGZ’s N-terminus, containing two C2H2 zinc fingers, binds to tubulin, the mitotic kinase Aurora A, and microtubules, while BuGZ undergoes liquid-liquid phase separation through its intrinsically disordered C-terminal region. In combination, the respective N- and C-terminal domain characteristics lead to enrichment of tubulin and Aurora A in BuGZ condensates, which in turn promote microtubule polymerization and Aurora A kinase activity, respectively (Huang et al., 2017; Jiang et al., 2015). The multivalent binding of phase separated BuGZ also leads to microtubule bundling (Jiang et al., 2015). BuGZ’s ability to promote microtubule polymerization by concentrating tubulin within BuGZ condensates is similar to a behavior observed in other phase-separating microtubule regulators, such as TPX2 and centrosome components, indicating that phase separation may play a role in mitotic spindle formation (King and Petry, 2020; Woodruff et al., 2017).

Through quantitatively analyzing BuGZ’s binding affinity to tubulin and microtubules, we found that BuGZ exhibits 10-fold higher affinity for GDP-tubulin than for GTP-tubulin, and a 210-fold higher affinity for GDP-tubulin than for microtubules. We further show that BuGZ promotes nucleotide exchange, converting GDP-tubulin to GTP-tubulin. We will discuss the implication of our findings on microtubule assembly dynamics.

## Results & Discussion

### BuGZ exhibits weak binding to microtubules

Previous studies have shown that BuGZ binds to microtubules and tubulin dimers and that BuGZ phase separation leads tubulin to concentrate inside the condensates, which in turn promotes microtubule polymerization. BuGZ droplets formed along microtubules also cause microtubule bundling (Jiang et al., 2015). However, there is an incomplete understanding of the interplay between BuGZ’s interactions with tubulin and microtubules given a lack of quantitative binding data. Therefore, we performed quantitative assays to determine the binding affinity between BuGZ and microtubules.

To measure equilibrium binding of BuGZ to microtubules, we used microtubules polymerized with a 1:10 mixture of biotin-labeled:unlabeled GTP-tubulin such that ≥1 biotin-tubulin is present per 12-protofilament turn, facilitating retrieval of the microtubules with paramagnetic streptavidin beads. Taxol was added to stabilize the polymerized microtubules. Given that taxol-stabilized microtubules remain intact at low temperatures, we performed all equilibrium binding experiments at 4 °C to suppress BuGZ’s tendency to undergo phase separation (Schiff et al., 1979; Jiang et al., 2015). Under these conditions, we mixed varying concentrations of taxol-stabilized microtubules (0 µM, 5.5 µM, 11 µM, 22 µM, 44 µM, 110 µM) with streptavidin-coated paramagnetic beads. Purified *X.laevis* BuGZ was added to a final concentration of 1.19 µM and the reaction was allowed to reach equilibrium. Then, microtubule-bound BuGZ was pulled down by a magnet. The supernatant was collected and BuGZ depletion was measured by quantitative immunoblotting. The fraction of BuGZ bound to microtubules was plotted against microtubule concentrations and then fitted with a rectangular hyperbola (see Materials and Methods). The equilibrium dissociation constant (K_D_) was determined from the fit. The K_D_ of BuGZ for taxol-stabilized microtubules is 9.45 µM (95% CI: 7.72 µM - 11.50 µM) (Figure 1A). Since a similar fraction of BuGZ was bound to 10 µM unsheared or sheared microtubules, BuGZ does not show an obvious preference to microtubule ends or the lattice under our assay conditions (Figure 1A, 1B).

**Figure 1.**
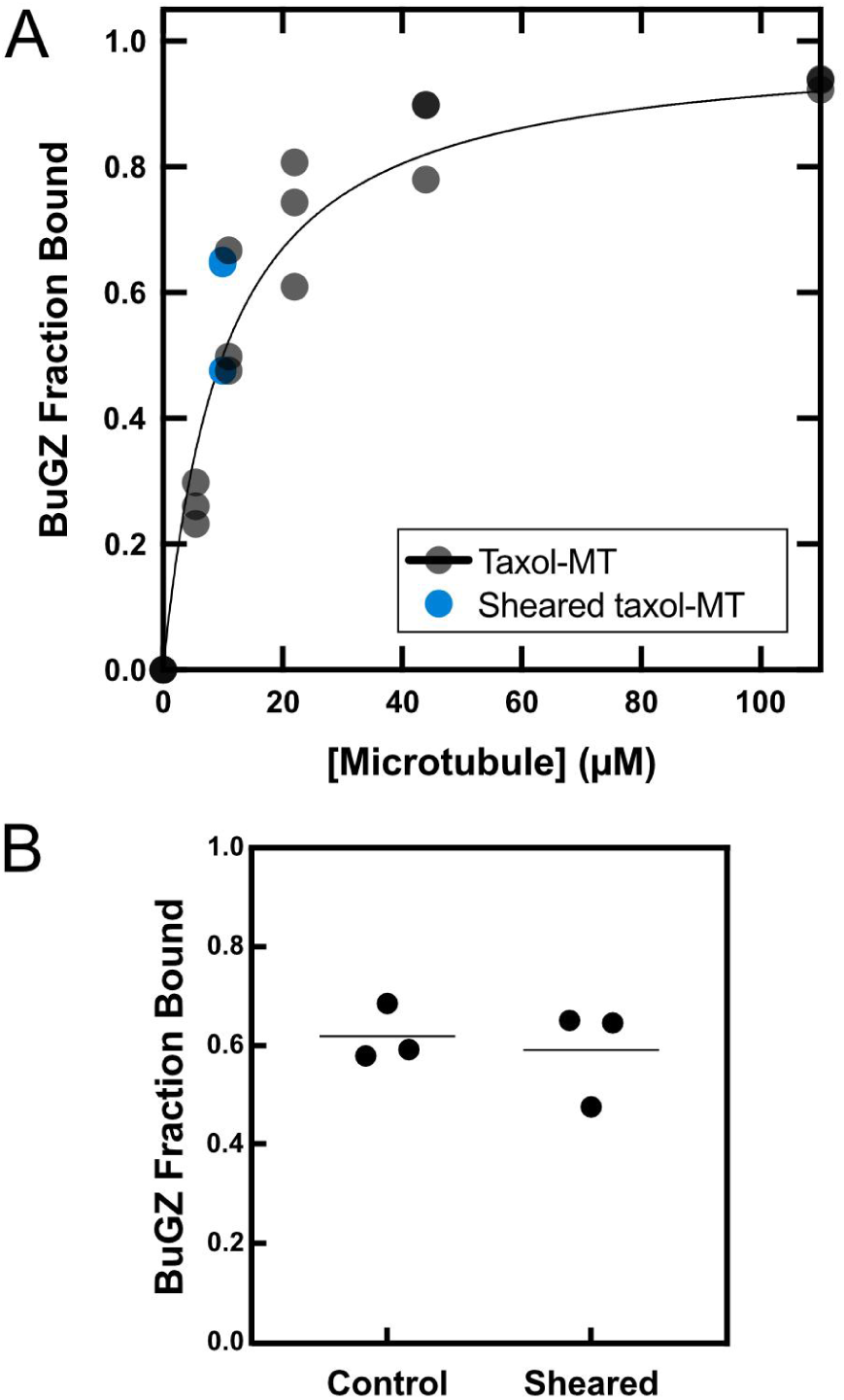
BuGZ exhibits weak binding to microtubules. (A) BuGZ binding isotherm to taxol-microtubules. K_D_ = 9.45 µM (95% CI = 7.72 µM-11.50 µM). Black data points correspond to the BuGZ fraction bound to taxol-MT. Curve-fit determined by *Equation 1* (Materials and Methods). Data from (B) for the BuGZ fraction bound to 10 µM sheared taxol-MT shown in blue. (B) BuGZ fraction bound to 10 µM taxol-microtubules and sheared taxol-microtubules. Data shown with the mean of three replicates.

### BuGZ preferentially binds GDP-tubulin over GTP-tubulin

The findings above show that the affinity of BuGZ for microtubules is relatively weak compared to other microtubule associated proteins whose binding affinities were previously reported (Folker et al., 2005; Spittle et al., 2000; Butner and Kirschner, 1991; Seeger and Rice, 2010). Since BuGZ also binds to tubulin and BuGZ condensates concentrate tubulin dimers, we next measured how well BuGZ binds free tubulin dimers (Jiang et al., 2015). We first investigated BuGZ’s binding affinity for GTP-tubulin. Streptavidin-coated paramagnetic beads were incubated with varying concentrations (0 µM, 0.1 µM, 1 µM, 2 µM, 5 µM, 10 µM, 20 µM) of biotinylated GTP-tubulin and then BuGZ was added to a final concentration of 1.19 µM. The reaction was performed at 4 °C in the presence of nocodazole to prevent the assembly of microtubules. The beads were retrieved via magnetic separation, and BuGZ equilibrium binding was assessed by measuring BuGZ depletion from the supernatant via quantitative immunoblotting. The K_D_ for BuGZ binding to GTP-tubulin was determined to be 477 nM (95% CI: 275.1 nM - 757.5 nM) (Figure 2A). The 20-fold stronger binding affinity of BuGZ to GTP-tubulin than to microtubules prompted us to measure the binding affinity of BuGZ to GDP-tubulin using the same assay. We found that the K_D_ for BuGZ binding to GDP-tubulin was 45.3 nM (95% CI: 21.7 nM - 78.5 nM), representing a 10-fold stronger affinity than for GTP-tubulin and 210-fold stronger affinity than for taxol-microtubules (Figure 2B). Control experiments using mEGFP in solution with bead-bound tubulin show no differential effect in GDP or GTP conditions (Figure 2C).

**Figure 2.**
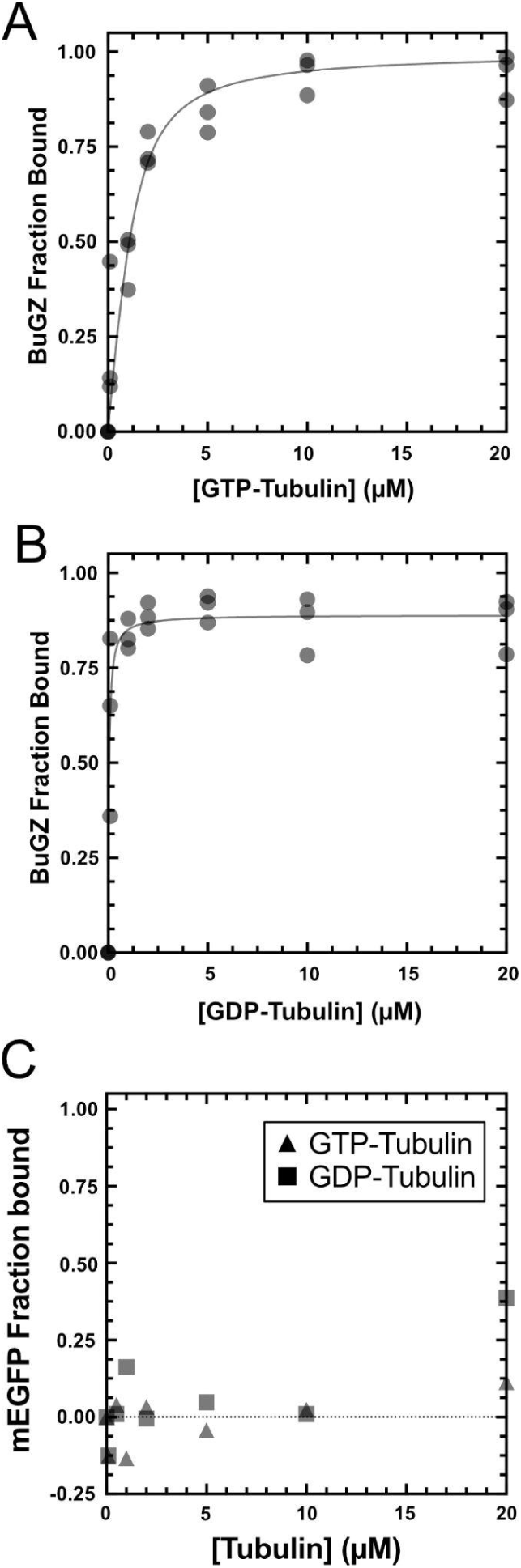
BuGZ exhibits preferential binding to GDP-tubulin over GTP-tubulin. (A) BuGZ binding isotherm to GTP-tubulin. K_D_ = 476.6 nM (95%CI = 275.1 nM - 757.5 nM). Curve-fit determined by *Equation 1* (Materials and Methods). (B) BuGZ binding isotherm to GDP-tubulin. K_D_ = 45.3 nM (95% CI = 21.7 nM - 78.5 nM). Curve-fit determined by *Equation 2* (Materials and Methods). (C) mEGFP as a binding assay control does not exhibit differential behavior with GDP-tubulin or GTP-tubulin. Bound mEGFP with increasing tubulin concentrations: 0 µM, 0.1 µM, 0.5 µM, 1 µM, 2 µM, 5 µM, 10 µM, 20 µM. Binding is not affected by tubulin, GDP, nor GTP. Circles denote GTP-tubulin binding data. Squares denote GDP-tubulin binding data.

### The N-terminal zinc finger-containing domain is responsible for the preferential binding of BuGZ to GDP-tubulin

To further investigate the ability of BuGZ to recognize the nucleotide-state of tubulin, we used AlphaFold to generate a predicted protein structure of BuGZ (Fig. 3A) (Jumper et al., 2021). Since most of the sequence following the N-terminal zinc finger motifs of BuGZ is predicted to be intrinsically disordered, the structure prediction beyond the N-terminal region is considered low confidence by AlphaFold (Fig. 3A). Next, we used the multimer model of AlphaFold to predict the structure of a BuGZ in-complex with a tubulin dimer (Fig. 3B). AlphaFold correctly predicted the tubulin dimer structure as previously determined by cryo-electron microscopy (Nogales et al., 1998b). AlphaFold also predicted that the C-terminal GLEBS motif of BuGZ, containing a short α-helix, is positioned close to the interface of α-tubulin and β-tubulin implying that the GLEBS domain could interact with tubulin (Fig 3B). Consistent with our previously reported finding, AlphaFold also predicted an interaction between the N-terminus of BuGZ and the tubulin dimer (Jiang et al., 2014, 2015). Based on these analyses, we constructed and purified two mutants of BuGZ: BuGZ-ΔGLEBS with a deletion of the 32 amino acids containing the GLEBS motif in the Proline-Rich Region 2 (PRR2) and the N-terminal 92 amino acids of BuGZ (BuGZ-NTD), containing the two zinc fingers (Figure 3C).

**Figure 3.**
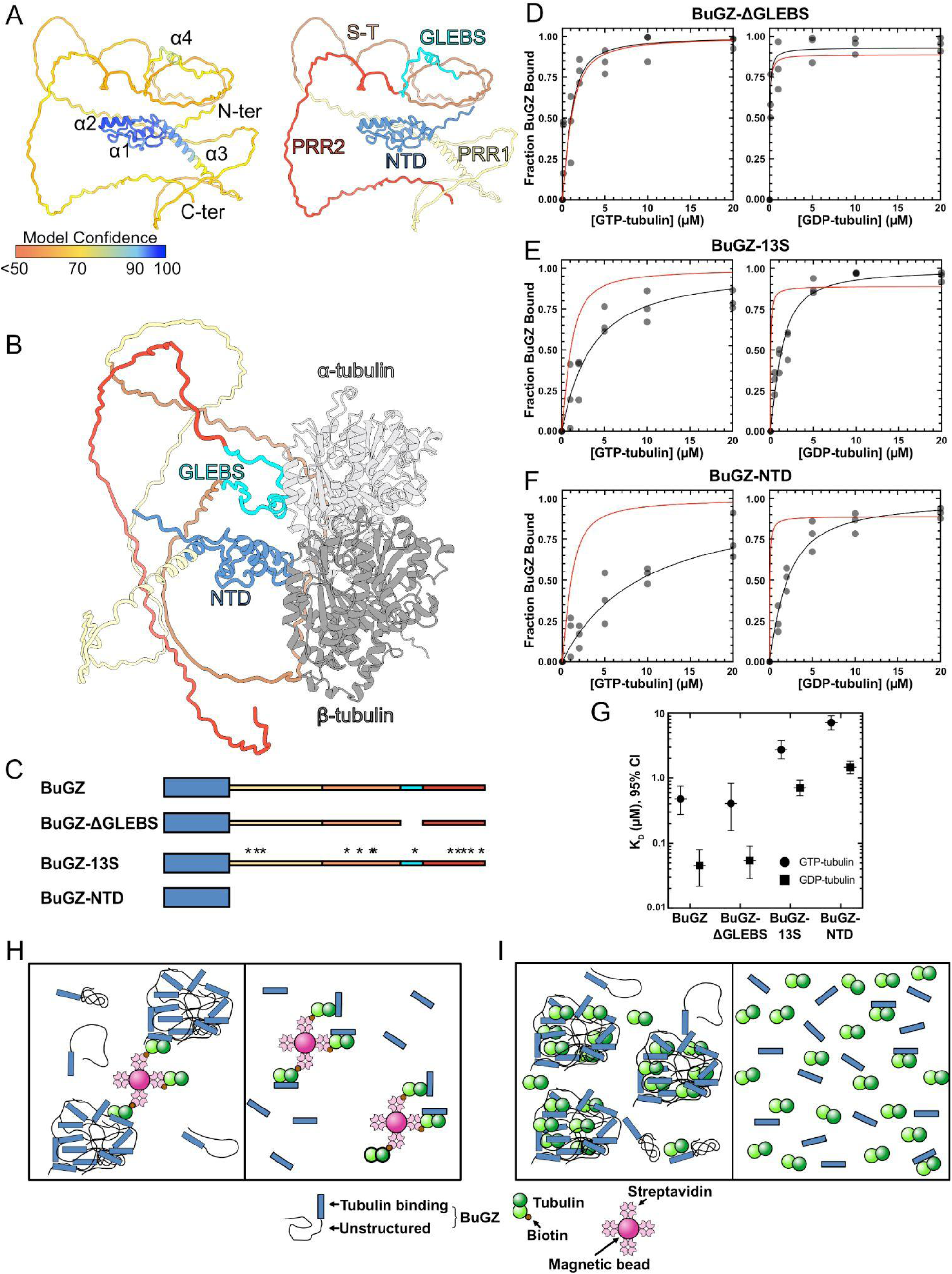
BuGZ’s N-terminal domain is responsible for nucleotide-specific binding of tubulin and loss of BuGZ phase separation results in weaker tubulin binding. (A) AlphaFold structure prediction for *X.laevis* BuGZ. Left: color-coded by Model Confidence score. Dark blue, Very high (> 90). Light blue, High (90 - 70). Yellow, Low (70 - 50). Orange, Very low (< 50). Right: Color-coded by protein region. Dark blue, N-terminal domain (NTD). Yellow, proline-rich region 1 (PRR1). Orange, serine-threonine rich region (S-T). Cyan, GLEBS domain (GLEBS). Red, proline-rich region 2 (PRR2). (B) AlphaFold structure prediction for BuGZ in-complex with α-tubulin and β-tubulin. BuGZ color-coding is the same as (A, right). α-tubulin, light gray. β-tubulin, dark gray. (C) Schematics of BuGZ constructs. BuGZ (top), BuGZ-ΔGLEBS (middle-top), BuGZ-13S (middle-bottom), BuGZ-NTD (bottom). Domain color-coding is the same as (A, right). Asterisks indicate the 13 Ser substitutions in BuGZ-13S. (D) Left, BuGZ-ΔGLEBS binding isotherm to GTP-tubulin. K_D_ = 408.1 nM (95% CI = 156.3 nM-834.4 nM). Right, BuGZ-ΔGLEBS binding isotherm to GDP-tubulin. K_D_ = 54.2 nM (95% CI = 28.6 nM-90.6 nM). (E) Left, BuGZ-13S binding isotherm to GTP-tubulin. K_D_ = 2.75 μM (95% CI = 1.97 μM-3.78 μM). Right, BuGZ-13S binding isotherm to GDP-tubulin. K_D_ = 710 nM (95% CI = 533 nM-926 nM). (F) Left, BuGZ-NTD binding isotherm to GTP-tubulin. K_D_ = 7.14 μM (95% CI = 5.53 μM-9.17 μM). Right, BuGZ-NTD binding isotherm to GDP-tubulin. K_D_ = 1.47 μM (95% CI = 1.18 μM-1.82 μM). Note: Red lines in panels D, E, and F show the binding curve of full-length wild-type BuGZ from Figure 2. Curve-fits for (D, E, F) determined by Equations 1 and 2 (Materials and Methods). (G) Equilibrium binding constants (K_D_) for BuGZ and BuGZ mutants shown in μM with error bars as 95% confidence intervals. Circles denote GTP-tubulin-related data points and squares denote GDP-tubulin-related data points. (H-I) BuGZ interactions with tubulin dimers. (H) Cartoon schematic of BuGZ oligomer (left) or BuGZ-NTD monomer (right) binding to bead-bound tubulin. (I) Cartoon schematic of unrestricted binding in solution of BuGZ (left) or BuGZ-NTD (right) to tubulin.

Using the same binding assay as described above, we measured the K_D_ for the binding of each BuGZ mutant to GTP-tubulin and GDP-tubulin. We found that BuGZ-ΔGLEBS binds to GTP-tubulin and GDP-tubulin at a K_D_ of 408.1 nM (95% CI: 156.3 nM-834.4 nM) and 54.2 nM (95% CI: 28.6 nM-90.6 nM), respectively (Figure 3D). Since these affinities are nearly identical to those measured for wild-type BuGZ, we conclude that the GLEBS motif is not involved in BuGZ binding to tubulin. For BuGZ-NTD, the K_D_ for GTP-tubulin and GDP-tubulin binding was measured as 7.14 µM (95% CI: 5.53 µM-9.17 µM) and 1.47 µM (95% CI: 1.18 µM-1.82 µM), respectively (Figure 3F). This indicates that BuGZ’s N-terminal 92 amino acids containing the two zinc fingers exhibit preferential binding for GDP-tubulin binding over GTP-tubulin.

The reduced binding affinity of BuGZ-NTD to tubulin, relative to full-length BuGZ, indicates that BuGZ’s C-terminal intrinsically disordered region plays a role in BuGZ-tubulin interactions. To investigate this finding further, we performed binding assays using a previously reported BuGZ mutant, BuGZ-13S (Jiang et al., 2015). This mutant is identical to wild-type BuGZ with the exception of 13 aromatic amino acid residues (phenylalanine or tyrosine) mutated to serine. While retaining BuGZ’s C-terminal intrinsically disordered region, the loss of aromatic residues results in greatly reduced phase separation of BuGZ (Jiang et al., 2015). We found that the K_D_ for BuGZ-13S binding to GTP-tubulin and GDP-tubulin are 2.75 µM (95% CI: 1.97 µM-3.78 µM) and 710 nM (95% CI: 533 nM-926 nM), respectively (Figure 3E). The stronger binding affinity of BuGZ-13S to GTP- and GDP-tubulin, relative to BuGZ-NTD, shows that the C-terminal intrinsically disordered region contributes to BuGZ-tubulin binding with the N-terminal domain of BuGZ contributing to the interaction bias toward GDP-tubulin.

Proteins that form condensates exhibit the propensity to form small oligomers even under conditions that disfavor phase separation (Chattaraj et al., 2024; Martin et al., 2021). Although our quantitative measurements of interaction affinities are performed at 4 °C, which suppresses the formation of BuGZ condensates visible under light microscopy, small BuGZ oligomers may still form. These oligomers would lead to an increased depletion of BuGZ by the biotinylated tubulin bound on beads in our assay, which could explain why the phase separation property of BuGZ contributes to increased tubulin binding (Figure 3H-I).

### BuGZ promotes GTP exchange into GDP-tubulin

Nucleotide dissociation from tubulin is known to be rapid (Brylawski and Caplow, 1983). Since cells have a 10-fold higher concentration of GTP over GDP, it is thought that most GDP-tubulin produced upon microtubule disassembly are rapidly converted into GTP-tubulin (Traut, 1994). However, in cellular states with increased microtubule polymerization and depolymerization, such as in mitosis where microtubule turnover has been measured to be 18-fold faster than in interphase, tubulin may require a nucleotide exchange factor to sustain the higher levels of microtubule dynamicity as small perturbations can cause defects during spindle assembly and cell division (Saxton et al., 1984; Vicente and Wordeman, 2019). The ten-fold binding affinity bias of BuGZ to GDP-tubulin over GTP-tubulin prompted us to investigate whether such preferential binding could facilitate the conversion of GDP-tubulin to GTP-tubulin. To test this possibility, we compared the incorporation of GTP α-^32^P into GDP-tubulin in the presence of BuGZ at room temperature, a condition in which BuGZ would undergo increased phase separation relative to our binding assays performed at 4 °C. 1mM GTP including 3 nM GTP α-^32^P was added to a solution of 100 µM GDP-tubulin and 1mM GDP to achieve a 1:1 ratio of GTP:GDP. Then, BuGZ or BSA was added to a final concentration of 1.19 µM. We hypothesized that if BuGZ preferentially promotes nucleotide exchange on GDP-tubulin because of its stronger binding affinity, the presence of BuGZ would result in higher GTP incorporation into tubulin, even in the presence of a 1:1 ratio of available GTP and GDP. At equilibrium, nucleotides were cross-linked to tubulin via ultraviolet radiation and unbound nucleotides were removed by size-exclusion chromatography. The eluate containing tubulin and BuGZ was assessed by scintillation counting (Figure 4A). In the presence of BuGZ, GTP incorporation, measured by the exchange of GDP for ^32^P-labeled GTP, increased by 63.9% (p=0.03) relative to the control condition (Figure 4B). This shows that BuGZ acts as a tubulin GEF to promote GTP incorporation in tubulin.

**Figure 4.**
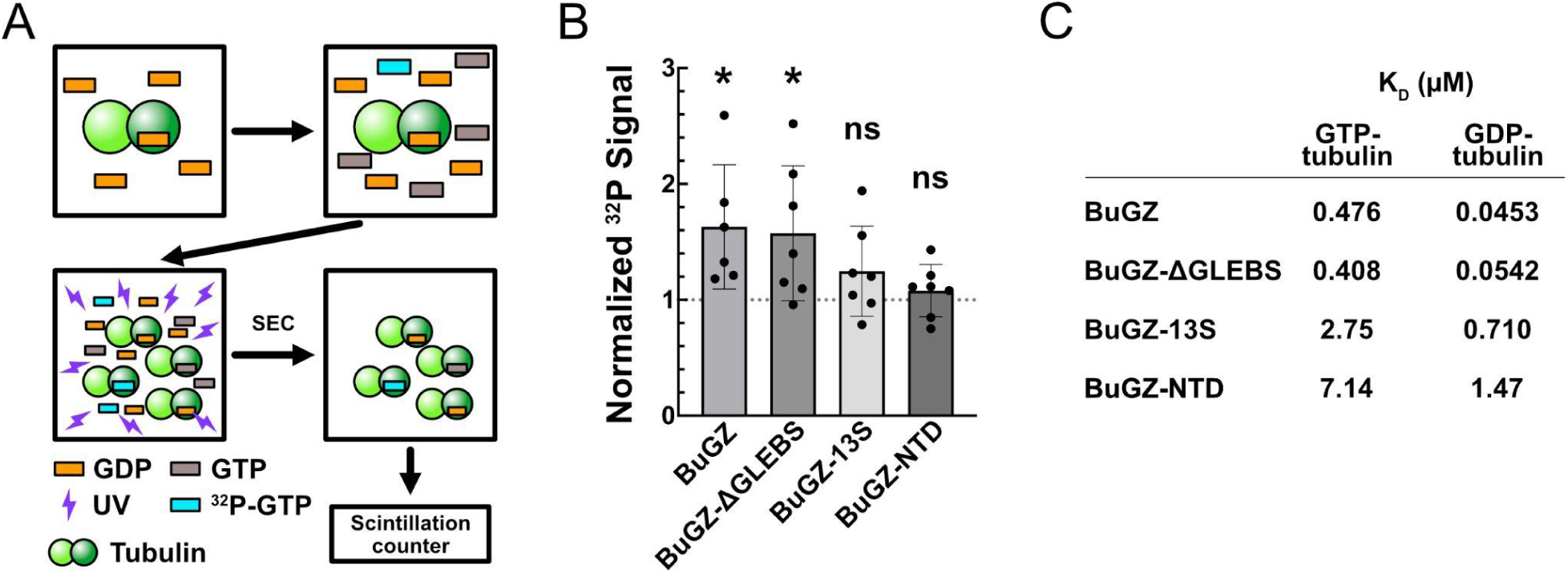
BuGZ acts as a guanine nucleotide exchange factor (GEF) for tubulin. (A) Workflow of BuGZ GEF assay. Tubulin dimers in 1mM GDP buffer are supplemented with nocodazole, BuGZ (or BSA control), equimolar GDP and GTP (the GTP includes the spike-in of ^32^P-GTP). After 15 minute incubation, the mixture was subjected to UV-mediated crosslinking and then passed through a desalting column to remove unincorporated nucleotides. Then, incorporated ^32^P was measured by a scintillation counter. Light green = α-tubulin. Dark green = β-tubulin. Orange = GDP. Grey = GTP. Cyan = ^32^P -GTP. (B) Presence of BuGZ in solution with tubulin and equal molar quantities of GDP and GTP increases GTP incorporation by 63.9% relative to BSA control (p = 0.03). BuGZ-ΔGLEBS increases GTP incorporation by 57.4% (p = 0.03). BuGZ-13S increases GTP incorporation by 24.6% (ns). BuGZ-NTD increases GTP incorporation by 7.8% (ns). Mean and standard deviation shown in the bar graph. Wilcoxon signed-rank test: *p ≤ 0.05, ns, not significant, theoretical value = 1. (C) Summary table listing BuGZ and BuGZ mutant binding affinities for GTP- and GDP-tubulin in micromolar scale.

Next, we tested the three BuGZ mutants for nucleotide exchange activity using the same assay. At equilibrium, BuGZ-ΔGLEBS showed a significant increase of 57.4% (p=0.03) in GTP incorporation relative to controls (Figure 4B). Conversely, BuGZ-13S and BuGZ-NTD showed statistically insignificant increases of GTP incorporation by 24.6% (p>0.05) and 7.8% (p>0.05), respectively (Figure 4B). Therefore, the ability of BuGZ to promote GTP exchange of GDP-tubulin relies on both the N-terminal tubulin binding domain and BuGZ phase separation mediated by the unstructured C-terminal region.

## Discussion

Although extensive efforts have led to the discovery of many microtubule-associated proteins (MAP), there have been limited studies on their interactions with tubulin. There is also no reported effort on identifying a tubulin GEF, possibly because of the observed rapid dissociation of GDP or GTP from tubulin. Here, we show that BuGZ is a tubulin binding protein with 10-fold stronger affinity toward GDP-tubulin over GTP-tubulin and that it exhibits GEF activity toward tubulin. Studies have shown that some MAPs exhibit preferential binding to microtubule segments containing either GTP-tubulin, such as those present at the plus-end of polymerizing microtubules, or GDP-tubulin found in the microtubule lattice (Goodson and Jonasson, 2018). Although these studies suggest that MAPs may preferentially bind to GTP- or GDP-bound tubulin, nucleotide-specific MAP-tubulin interactions have only been reported for two MAPs, CLIP-170 and XMAP215 (Ayaz et al., 2012; Folker et al., 2005). In contrast to BuGZ which binds to GDP-tubulin at K_D_ = 45.3 nM and GTP-tubulin at K_D_ = 477 nM mediated by its zinc finger-containing N-terminal domain (Figure 2A-B, 5C), CLIP-170 and a subdomain of XMAP215 called TOG exhibit similar binding affinity to both GDP- and GTP-tubulin at K_D_ ≈ 45 nM and K_D_ ≈ 235 nM, respectively (Folker et al., 2005; Ayaz et al., 2012). Our finding of BuGZ as the first tubulin binding protein that shows preferential interactions with GDP-tubulin over GTP-tubulin reveals the need to further quantitatively study the ability of various MAPs to bind to tubulin bound to GDP or GTP (Gache et al., 2005; Yu et al., 2016). Further characterization in this manner may uncover an underappreciated regulatory role of MAPs in modulating microtubule assembly and function in different cellular contexts.

Our finding of BuGZ as a tubulin GEF is unexpected because of the prevailing idea that there is an abundance of intracellular GTP and that the GTP:GDP ratio is high in cells (Traut, 1994). Coupled with the rapid intrinsic nucleotide dissociation from tubulin should allow GDP-tubulin to quickly exchange into GTP-tubulin (Brylawski and Caplow, 1983). However, a recent study shows that the availability of GTP in the cell is rate-limiting in nucleocytoplasmic transport, indicating that GTP may become limiting when energy demand is high (Scott et al., 2024). Mitosis is an energy demanding process as it involves structural reorganization of the whole cell. Moreover, studies have shown that microtubule growth is diffusion-limited such that increasing the density of growing microtubule tips and increasing the local concentration of GTP-tubulin dimers causes a decrease and increase in microtubule growth, respectively (Geisterfer et al., 2020; Odde, 1997; Geel et al., 2020). In mitosis, the rapid growth and shrinkage of microtubule plus and minus ends, respectively, in the spindle could locally deplete GTP-tubulin. By binding to the GDP-tubulin generated from spindle microtubule depolymerization, BuGZ could increase the rate of GTP-tubulin production and subsequently, the 10-fold lower affinity for GTP-tubulin would mediate BuGZ’s release of the tubulin, post-exchange, to support the highly dynamic spindle microtubules. How BuGZ functions as a GEF remains unclear, but its preferential binding to GDP-tubulin suggests that it may not facilitate equal nucleotide exchange of GDP- and GTP-tubulin as seen for other GEFs but instead specifically facilitates nucleotide exchange GDP-tubulin. Since the zinc finger domain of BuGZ can be produced in bacteria with high quantity and purity, it should be possible to solve the structure of the BuGZ-tubulin complex, which should shed light on how BuGZ functions as a tubulin GEF.

## Materials and Methods

### Cloning, protein expression, protein purification

*Xenopus laevis* BuGZ cDNA, codon-optimized for *Spodoptera frugiperda* Sf9 expression, was cloned into a baculovirus vector (Gibco pFastBac Dual Expression Vector) via Gibson Assembly (NEB Gibson Assembly Master Mix). In the assembled plasmid, BuGZ (NCBI: NM_001086855.1) is tagged on its amino terminus with: four leader amino acids, 6x His tag, GS linker, and TEV protease cleavage site (full tag sequence:

MSYY-HHHHHH-GSG_4_SG_4_S-ENLYFQG). Vectors were then transfected into Sf9 cells (Gibco Sf9 cells in Sf-900 III SFM), using Gibco Cellfectin II Reagent according to ThermoFisher Bac-to-Bac Baculovirus Expression System instructions, to generate P0 viral stock. After 3 rounds of viral amplification, the resulting P3 virus was used to infect Sf9 cells for protein expression. 72 hrs post-infection, cells were collected into 100 mL pellets via centrifugation at 1,000 g. Pellets were snap-frozen in liquid nitrogen and stored at -80 °C. For purification, pellets were thawed and resuspended in lysis buffer (LyB): 20 mM KH_2_PO_4_, 500 mM NaCl, 25 mM Imidazole, 1 mM MgCl_2_, 1 mM β-mercaptoethanol, 2.5% Glycerol, 0.01% Triton X-100, Roche cOmplete, EDTA-free Protease Inhibitor Cocktail, pH 7.4. The cell suspension was sonicated using Misonix Sonicator 3000 (2 min total process time, 30 s on, 30 s off, power level 2.0) on ice and then clarified by centrifugation at 15,000 x g for 30 min. Clarified lysate was then filtered through a MilliporeSigma Millex-GP 0.22 μm PES membrane filter unit. Lysate was loaded onto a 1 mL HisTrap HP column (Cytiva) pre-equilibrated in Buffer A (same as LyB without protease inhibitor cocktail) and ran on the FPLC (Cytiva Äkta pure). The column was then washed with 1X lysate volume of 85% Buffer A/15% Buffer B (20 mM KH_2_PO_4_, 150 mM NaCl, 300 mM Imidazole, 1 mM MgCl_2_, 1 mM β-mercaptoethanol, 2.5% Glycerol, 0.01% Triton X-100, pH 7.4). Bound protein was eluted with a 30 mL linear gradient from 15% to 100% Buffer B.

BuGZ-containing fractions were identified by running fractions via SDS-PAGE and staining with Invitrogen SimplyBlue SafeStain (Invitrogen LC6060). BuGZ-positive fractions were pooled and concentrated to < 500 μL using Millipore Amicon Ultra centrifugation units with 30 kDa MW cut off (10 kDa MW cut-off for BuGZ-NTD). Contaminating proteins were removed from the concentrated BuGZ protein solution by gel filtration using a Superdex 200 Increase 10/300 GL column equilibrated in Buffer C (80 mM PIPES pH 6.8, 100 mM KCl, 1 mM MgCl_2_, 50 mM Sucrose, 1 mM EGTA). Fractions of the eluate containing BuGZ were again identified via SDS-PAGE and were then pooled, and concentrated using Millipore Amicon Ultra centrifugation units, and snap-frozen in liquid nitrogen for storage at -80 °C. The purity and concentration of BuGZ was determined by comparing in-gel SimplyBlue SafeStain staining to staining of known amounts of bovine serum albumin.

### Measurement of BuGZ binding to Taxol-stabilized microtubules

10 mg/mL tubulin (Cytoskeleton T240) and 1 mg/mL biotin-tubulin (Cytoskeleton T333P) were mixed in equal volumes in a buffer containing BRB80 (80mM PIPES, 1mM MgCl_2_, 1mM EGTA, pH 6.8) + 1 mM GTP. The tubulin solution was diluted 1:1 in a solution containing BRB80, 2 mM DTT, 2 mM GTP, 20 μM taxol (taxol buffer 1) and incubated for 20 min at 37 °C. Pierce Streptavidin Magnetic Beads (ThermoFisher Scientific 88816) were equilibrated in a buffer containing BRB80, 1 mM DTT, 1 mM GTP, 10 μM taxol (taxol buffer 2). The microtubule assembly mixture was added to 20 μL streptavidin-coated bead slurry and incubated for 20 min at 4 °C, then washed 3 times with taxol buffer 2. After the final wash, the beads were resuspended in a 10 μL solution containing taxol buffer 2 and 1.19 μM BuGZ. After 20 min incubation at 4 °C, the beads were collected with a magnet and the supernatant was removed, and mixed with an equal volume of 2X SDS sample buffer for immunoblot analysis. For experiments with sheared microtubules, the taxol-stabilized microtubules were passed through a 30 G syringe needle 10 times immediately prior to addition to streptavidin-coated beads.

### Measurement of BuGZ binding to GTP- and GDP-tubulin

Varying volumes of solution containing 1 mg/mL (∼10 μM) biotin-tubulin, 200 μM nocodazole, 1 mM GTP or GDP, and BRB80 were added to Pierce Streptavidin Magnetic Beads equilibrated in the same buffer to decorate them with the desired amount of tubulin. After 20 min. incubation at 4 °C, beads were washed 3 times in the same buffer and the supernatant was removed. The magnetic beads, now decorated with varying amounts of biotin tubulin, were resuspended in equal volumes of a solution containing 1.19 μM BuGZ, 200 μM nocodazole, 1 mM GTP or GDP, and BRB80. After 15 min incubation at 4 °C, beads were collected with a magnet and supernatant was retained (mixed 50% v/v with 2X Laemmli buffer) for immunoblot analysis.

### Gel electrophoresis and immunoblot analysis

Supernatants from binding experiments were boiled at 95 °C for 10 min and spun down in a benchtop centrifuge at 10,000 g for 3 min. Samples were loaded into an SDS polyacrylamide gel and separated by electrophoresis in a Tris-Glycine buffer (25 mM Tris, 250 mM glycine, 3.5 mM SDS). Following electrophoresis, proteins were transferred onto nitrocellulose membrane (Cytiva Amersham 10600002) for 2 hrs at 4 °C in transfer buffer (50 mM Tris, 125 mM glycine, 3.5 mM SDS, 20% methanol). After transfer, membranes were incubated in a block buffer (5% w/v skim milk, Tris-buffered saline pH 7.4) for 1 hr at room temperature, then probed with primary antibodies to the 6X His tag (for BuGZ recognition) in 5% w/v skim milk, Tris-buffered saline, 0.02% Tween-20, for 1 hr at room temperature. After washing 3 times for 10 min each in tris-buffered saline, membranes were incubated in a secondary antibody solution (Tris-buffered saline, 0.02% Tween-20, antibody) for 1 hr at room temperature. After washing 3 times for 10 min in tris-buffered saline, membranes were imaged in a LI-COR CLx system. Anti-6X His tag antibody (Abcam ab18184) was diluted 1:1000, and the secondary antibody, LI-COR IRDye 680RD Goat anti-Mouse IgG (926-68070), was diluted 1:10,000.

### Nucleotide exchange assay

A solution containing 100 μM tubulin, 1 mM GDP, BRB80 was mixed with a 1/10 volume of 10X nucleotide exchange buffer (10 mM GTP, 33.3 nM GTP α-^32^P (PerkinElmer BLU506H250UC), 1 mM nocodazole, BRB80) for a final solution containing 91 μM tubulin, 0.9 μM GDP, 0.9 μM GTP, 2.7 nM GTP α-^32^P, 91 μM nocodazole, and BRB80. Immediately, a BRB80 solution containing either BuGZ or BSA was added to a final concentration of 1.19 μM of the protein. Corresponding controls were produced in the same manner except without the addition of tubulin. Mixtures were incubated at room temperature for 15 min. Then, samples were treated with UV for 5 min in a Stratagene UV Stratalinker 1800. Free nucleotides were removed with BioRad Micro Bio-Spin 6 gel columns (BioRad 7326222) equilibrated to BRB80 buffer according to manufacturer’s instructions. Flowthrough was mixed with a scintillation cocktail (RPI Bio-Safe II) and GTP α-^32^P was measured using PerkinElmer Tri-Carb 2810 TR. The measured signal from the no tubulin conditions for BSA or BuGZ were subtracted from its corresponding tubulin + BuGZ or BSA readouts. Then, the data were normalized as a ratio of the BuGZ to BSA ^32^P signals.

### Data analysis

All statistical analyses were performed using Graphpad Prism 10. Equilibrium dissociation constants were determined for conditions where the K_D_ was near or far exceeding the total concentration of BuGZ (BuGZ to taxol-MT, BuGZ to GTP-tubulin, BuGZ-ΔGLEBS to GTP-tubulin, BuGZ-13S to GDP- and GTP-tubulin, BuGZ-NTD to GDP- and GTP-tubulin), using the function:

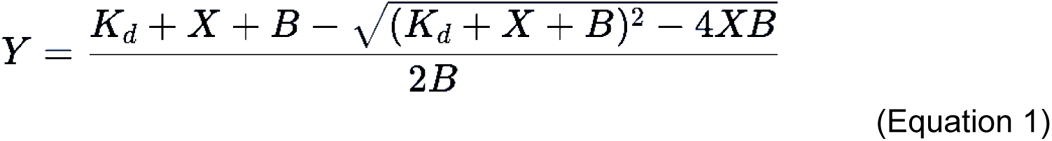

where X is the microtubule/tubulin concentration, B is the concentration of BuGZ, and Y is the BuGZ fraction bound.

For conditions where the K_D_ was far less than the total concentration of BuGZ (BuGZ to GDP-tubulin, BuGZ-ΔGLEBS to GDP-tubulin), equilibrium dissociation constants were determined using Prism 10’s ‘Hyperbola’ function:

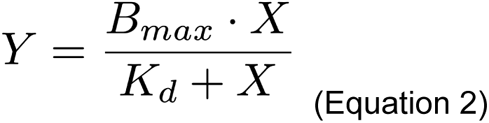

where Bmax = 1, X is the concentration of BuGZ, and Y is the BuGZ fraction bound. Nucleotide exchange assays were analyzed using Wilcoxon signed-rank test against a hypothetical value of 1.

### Alphafold protein structure prediction

Protein structure predictions of BuGZ (UniProt: Q7ZXV8), and the complex of BuGZ, α-tubulin (Uniprot: Q71U36), and β-tubulin (Uniprot: Q9H4B7) were generated using a modified version of AlphaFold v2.3.2 on Google Colab (https://colab.research.google.com/github/deepmind/alphafold/blob/main/notebooks/AlphaFold.ipynb). Predicted structures were analyzed and color-coded using ChimeraX. Heat map legend corresponding to “Model Confidence” was generated using ChatGPT.

## Acknowledgments

The authors would like to thank Ashish Tiwary (Carnegie Institution for Science) for helpful discussion and Kathryn A. Diederichs for assistance with protein structure prediction visualization. The study was funded by NIH NIGMS GM106023 (awarded to Yixian Zheng and Robert Goldman), NIH NIGMS GM110151 (awarded to Yixian Zheng), NIH NIGMS F32 GM142145 (awarded to Ross T.A. Pedersen), and NIH NIGMS R35 GM122569 (awarded to Taekjip Ha).

